# Harmonization of cortical thickness measurements across scanners and sites

**DOI:** 10.1101/148502

**Authors:** Jean-Philippe Fortin, Nicholas Cullen, Yvette I. Sheline, Warren D. Taylor, Irem Aselcioglu, Phil Adams, Crystal Cooper, Maurizio Fava, Patrick J. McGrath, Melvin McInnis, Ramin V. Parsey, Mary L. Phillips, Madhukar H. Trivedi, Myrna M. Weissman, Russell T. Shinohara

## Abstract

With the proliferation of multi-site neuroimaging studies, there is a greater need for handling non-biological variance introduced by differences in MRI scanners and acquisition protocols. Such unwanted sources of variation, which we refer to as “scanner effects”, can hinder the detection of imaging features associated with clinical covariates of interest and cause spurious findings. In this paper, we investigate scanner effects in two large multi-site studies on cortical thickness measurements, across a total of 11 scanners. We propose a set of general tools for visualizing and identifying scanner effects that are generalizable to other modalities. We then propose to use ComBat, a technique adopted from the genomics literature and recently applied to diffusion tensor imaging data, to combine and harmonize cortical thickness values across scanners. We show that ComBat removes unwanted sources of scan variability while simultaneously increasing the power and reproducibility of subsequent statistical analyses. We also show that ComBat is useful for combining imaging data with the goal of studying life-span trajectories in the brain.

## 1 Introduction

Large-scale efforts aimed at collecting diverse neuroimaging datasets for dissemination and sharing are rapidly growing in number and scale [Di Martino et al., 2014, Keator et al., 2013, Mennes et al., 2013]. Having multiple scan sites is necessary in large-scale studies due to logistical issues and geographic variability in subject populations [Van Horn and Toga, 2009]. However, a major drawback of combining neuroimaging studies across sites is the introduction of non-biological sources of variability to the data, typically related to the image acquisition protocol and hardware.

Properties of MRI scanners such as field strength, manufacturer, gradient nonlinearity, subject positioning, and longitudinal drift have been long understood to increase bias and variance in the measurement of brain volume changes [Takao et al., 2011], regional cortical thickness [Han et al., 2006], voxel-based morphometry [Takao et al., 2014], and structural, functional, and diffusion images in general [Jovicich et al., 2006, Takao et al., 2011]. Such unwanted sources of bias and variability are typically included as confound variables in the analysis of neuroimaging data. Recent work has suggested that standard methods for including confound variables for the prediction of an outcome using neuroimaging data perform no better than baseline models which ignore confounding [Rao et al., 2017]. Furthermore, non-biological confounders typically have a *priori* unpredictable effects, thus compromising consistency and reproducibility of the downstream analyses across studies. This suggests that non-biological sources of variability should be handled differently. Similar to *batch effects* in genomics (see Leek et al. [2010] for a review of batch effects), we use the term *scanner effects* in neuroimaging to refer to unwanted variation that is (1) non-biological in nature and (2) associated with differential scanning equipment or parameter configurations. Because different imaging sites use different physical scanners, site effects are one example of scanner effects.

Recently, ComBat [Johnson et al., 2007], a batch-effect correction tool commonly used in genomics, has been adapted for the modeling and removal of site effects in multi-site DTI studies [Fortin et al., 2017]. ComBat was found to be an effective harmonization technique that both removes unwanted variation associated with site and preserves biological associations in the data.

In this paper, we propose to use ComBat for harmonizing cortical thickness measurements obtained from multiple sites. We investigate this in region-level cortical thickness measurements in two large multi-site datasets: the Establishing Moderators and Biosignatures of Antidepressant Response in Clinical care study (EMBARC) [Trivedi et al., 2016], a multi-center study with 4 sites, and the Vascular Depression: Longitudinal Changes (VDLC) study, which was conducted at Washington University in St-Louis and Duke University, and used a total of 7 scanners. We first propose a set of general tools for the visualization and identification of site effects that are generalizable to other modalities. We then harmonize the data using ComBat, and compare to two other harmonization methods: residuals and phenotype-adjusted residuals. We show that Combat is successful at removing scanner and site effects, while preserving the variability associated with biology. We also show that ComBat can be used to combine datasets across multiple sites for the study of life-span trajectories.

**Table 1.** Description of the EMBARC and Vascular study samples

|  | EMBARC | Vascular |
| --- | --- | --- |
| N sites/scanners | 4 | 7 |
| N subjects | 187 | 236 |
| N females (%) | 116 (62) | 139 (59) |
| Age range | [18,65] | [58,95] |

## 2 Methods

### 2.1 Data and Preprocessing

#### EMBARC dataset

The EMBARC study aims to identify moderators and mediators of antidepressant response in adult patients with Major Depressive Disorder [Trivedi et al., 2016, Webb et al., 2016]. The dataset includes structural images, demographic variables and clinical variables. Participants were 200 unmedicated depressed individuals with Major depressive disorder and 40 healthy individuals recruited for EMBARC. Subjects were 18-65 years old, had to report age of depression onset before age 30 and had to be fluent in English. Clinical variables included the Hamilton Depression Rating Scale (HAMD) [Hamilton, 1960], the Mood and Anxiety Symptom Questionnaire (MASQ) [Watson and Clark, 1991], the Snaith-Hamilton Pleasure Scale [Snaith et al., 1995], the Spielberger State-Trait Anxiety Inventory (STAI) [Spielberger, 1983] and the Quick Inventory for Depression Symptomatology (QIDS) depression score [Rush et al., 2003].

The scans were acquired at four different imaging sites, with acquisition protocols described in Greenberg et al. [2015]. The four sites were Columbia University (CU), University of Texas South-western (TX), Massachusetts General Hospital (MG) and the University of Michigan (UM). All of the sites used 3T scanners, however the manufacturer differed from site to site: UM used a Philips Ingenia 3T scanner, TX used a Philips Achieva 3T scanner, MG used a Siemens TIM Trio 3T scanner and CU used a GE SIGNA HDx 3T scanner. Imaging parameters for each scanner are described in Greenberg et al. [2015]. After quality control, the final baseline dataset consisted of 187 subjects.

#### Vascular dataset

The Vascular Depression: Longitudinal Changes (VDLC) study aims to study the longitudinal effect of vascular disease in the pathogenesis of late-life depression (LLD). Participants were 117 individuals affected by LLD and 59 healthy controls, for a total of 236 participants. Participants were 58-95 years old. For the purpose of investigating site effects, we only considered one time point for each participant; we retained the scan from the last visit. Scans were acquired at two sites: Duke University and Washington University in St-Louis, across 7 different scanners, described in Table 2.

**Table 2.** Description of the scanners CU: Columbia University; MG: Massachusetts General Hospital; TX: University of Texas Southwestern; UM: University of Michigan; Duke: Duke University; WashU: Washington University in St-Louis.

|  | Location | Manufacturer | Platform | Field (T) | n | Label |
| --- | --- | --- | --- | --- | --- | --- |
| **EMBARC study** |  |  |  |  |  |  |
| Site 1 | CU | GE | SIGNA HDx | 3 | 46 | CU |
| Site 2 | MG | Siemens | TIM Trio | 3 | 26 | MG |
| Site 3 | TX | Philips | Achieva | 3 | 72 | TX |
| Site 4 | UM | Philips | Ingenia | 3 | 43 | UM |
| **Vascular study** |  |  |  |  |  |  |
| Scanner 1 | WashU | Siemens | Sonata | 1.5 | 23 | W_Sonata_A |
| Scanner 2 | WashU | Siemens | Sonata | 1.5 | 78 | W_Sonata_B |
| Scanner 3 | WashU | Siemens | TIM Trio | 3 | 16 | W_TIMTrio_A |
| Scanner 4 | WashU | Siemens | TIM Trio | 3 | 40 | W_TIMTrio_B |
| Scanner 5 | Duke | Siemens | TIM Trio | 3 | 24 | D_TIMTrio_A |
| Scanner 6 | Duke | Siemens | TIM Trio | 3 | 38 | D_TIMTrio_B |
| Scanner 7 | Duke | GE | SIGNA Excite | 1.5 | 17 | D_SIGNA |

### 2.2 Extraction of cortical thickness measurements

For the extraction of the cortical thickness measurements, we ran the ANTs cortical thickness pipeline, which has been shown to provide accurate and robust cortical thickness measurements [Tustison et al., 2014]. We used an average labeled template previously constructed from a subset of participants of the Open Access Series of Imaging Studies (OASIS) [Marcus et al., 2007] to identify the cortical regions for each subject of the EMBARC study.

### 2.3 Harmonization procedures

For the removal of site effects, we compare three different harmonization procedures: (1) Removal of site effects using linear regression without adjusting for biological covariates. We refer to the method as *Residuals*. (2) Removal of site effects using linear regression, adjusting for known covariates. We refer to the method as *Adjusted Residuals*. (3) Removal of site effects using *ComBat* [Johnson et al., 2007]. We also compare the three methods to the absence of harmonization, that we refer to as Raw. We describe below the different harmonization techniques.

For the different methods, we use the following notation. Let *y_ijv_
* be the *n* × 1 vector of cortical thickness measurement for imaging site *i*, for participant *j* and feature *v*, for a total of (*k* + 1) sites, *n* participants and *V* features. Depending on the cortical thickness modality, the features can either be ROIs, vertices or voxels. Furthermore, let **X** be the *p* × *n* matrix of biological covariates of interests, and let **Z** be the *k* × *n* matrix of site indicators (deviations from a baseline site).

#### 2.3.1 Residuals harmonization

The residuals harmonization method adjusts the images for site effects using linear regression. It does not take into account the potential confounding between the site variables and the biological covariates of interest in the study. The regression model can be written as

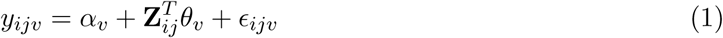

where *α_v_
* is the average cortical thickness for the reference site for feature *v* and where *θ_v_
* is the *k* × 1 vector of the coefficients associated with **Z** for feature *v*. We assume that the residual terms *ϵ_ijv_
* have mean 0. For each feature separately, we obtain an estimate 
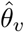
 of the parameter vector *θ_v_
* using regular ordinary least squares (OLS). The removal of site effects is done by subtracting the estimated site effects, that is we set the residuals-harmonized cortical thickness values to be

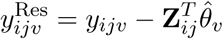

#### 2.3.2 Adjusted residuals harmonization

The adjusted residuals harmonization method supervises the removal of site effects by adjusting for biological covariates, using the following linear regression model:

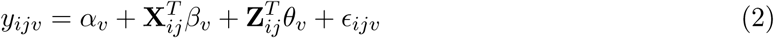

where *α_v_
* is the average cortical thickness for the reference site for feature *v*, where *θ_v_
* is the *k* × 1 vector of the coefficients associated with **Z** for feature *v* and where *β_v_
* is the *p* × 1 vector of coefficients associated with **X** for feature *v*. We assume that the residual terms *ϵ_ijv_
* have mean 0. For each feature separately, we obtain estimates and 
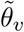
 and 
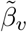
 using regular ordinary least squares (OLS) on the full model described in Equation 2. The removal of site effects is done by subtracting the estimated site effects only, that is we set the adjusted-residuals-harmonized cortical thickness values to be

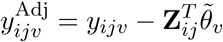

#### 2.3.3 ComBat harmonization

The Combat harmonization model [Johnson et al., 2007] extends the adjusted residuals harmonization model presented in Equation 2 in two ways: (1) it models site-specific scaling factors and (2) it uses empirical Bayes to improve the estimation of the site parameters for small sample sizes. It posits a unique linear model of location and scale at each feature, making the assumption that scanners (or sites) have both an additive and multiplicative effects on the data. The model assumes that the expected values of the imaging feature measurements can be modeled as a linear combination of the biological variables and the site effects, whose error term is modulated by additional site-specific scaling factors. The algorithm uses empirical Bayes to improve the estimation of the model parameters in small sample size studies. The ComBat model, originally developed for gene expression microarray data, was reformulated in Fortin et al. [2017] for the harmonization of DTI data scalar maps. Using the previous notation, the model can be written as

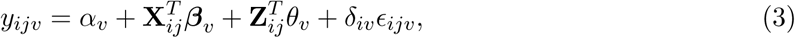

where *α_v_
* is the average cortical thickness for the reference site for feature *v*, where *θ_v_
* is the *k* × 1 vector of the coefficients associated with the site indicators Z for feature *v* and where *β_v_
* is the *p* × 1 vector of coefficients associated with X for feature *v*. We assume that the residual terms *ϵ_ijv_
* have mean 0. The parameters *δ_iv_
* descrive the multiplicative site effect of the *j*-th site on voxel *v*. Consistent with the ComBat model notation used in Fortin et al. [2017], we rewrite 
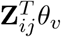
 as *γ_iv_*:

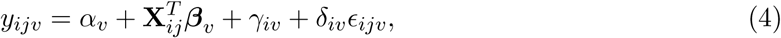

The procedure for the estimation of the site parameters *γ_iv_
* and *δ_iv_
* uses Empirical Bayes, and is described in Johnson et al. [2007] and Fortin et al. [2017]. The final ComBat-harmonized cortical thickness measusrements are defined as

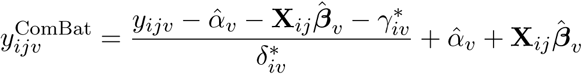

## 2.4 Methods evaluation framework

To investigate and correct site effects using ComBat, we performed a set of analysis tasks of increasing complexity on the cortical thickness data. We first performed an exploratory analysis to confirm the existence of site effects in the data. Next, we performed various univariate tests of significance to understand the relationships between individual features in the data and individual target variables. Finally, we applied various multivariate predictive models to understand how cortical thickness relates to target variables. Our analyses were aimed at both identifying and correcting site effects at multiple levels of complexity, along with understanding the specific effects of ComBat on downstream analysis.

## 3 Results

We first present visualization tools for investigating scanner effects in a multi-site study, and present several metrics to quantify such scanner effects. We use the cortical thickness measurements from the EMBARC study to illustrate the different methodologies. We next evaluate different harmonization procedures in the EMBARC study for the correction of site effects. We then show that the results are generalizable to other studies, using the VDLC dataset as an external validation. Finally, we combine and harmonize the EMBARC and VDLC studies, which have different age range, and show that it is possible to improve multi-site cross-sectional analyses of life-span trajectories by using ComBat harmonization.

### 3.1 Visualization of site effects

With high-dimensional data, routinely using fast and efficient visualization tools helps identifying biases and technical artifacts that affect the data globally. In this section, we present several visual tools, commonly used in genomics and other fields, for the identification and diagnostic of site effects in imaging data. We use the EMBARC dataset as an example dataset. We present the different diagnostic plots in Figure 1. First, for each subject, we summarize the cortical thickness measurements into a boxplot, and present the boxplots in Figure 1a in which the colors represent the four different imaging sites. We observe a global downwards shift in the cortical thickness measurements from the MG site, as well as increased variability in the measurements from the TX and UM sites relative to the two other sites. The four boxplots presented in Figure 1b summarize the distribution of the median cortical thicknesses at each site. This facilitates the visualization of the site-specific additive and scaling effects.

**Figure 1.**
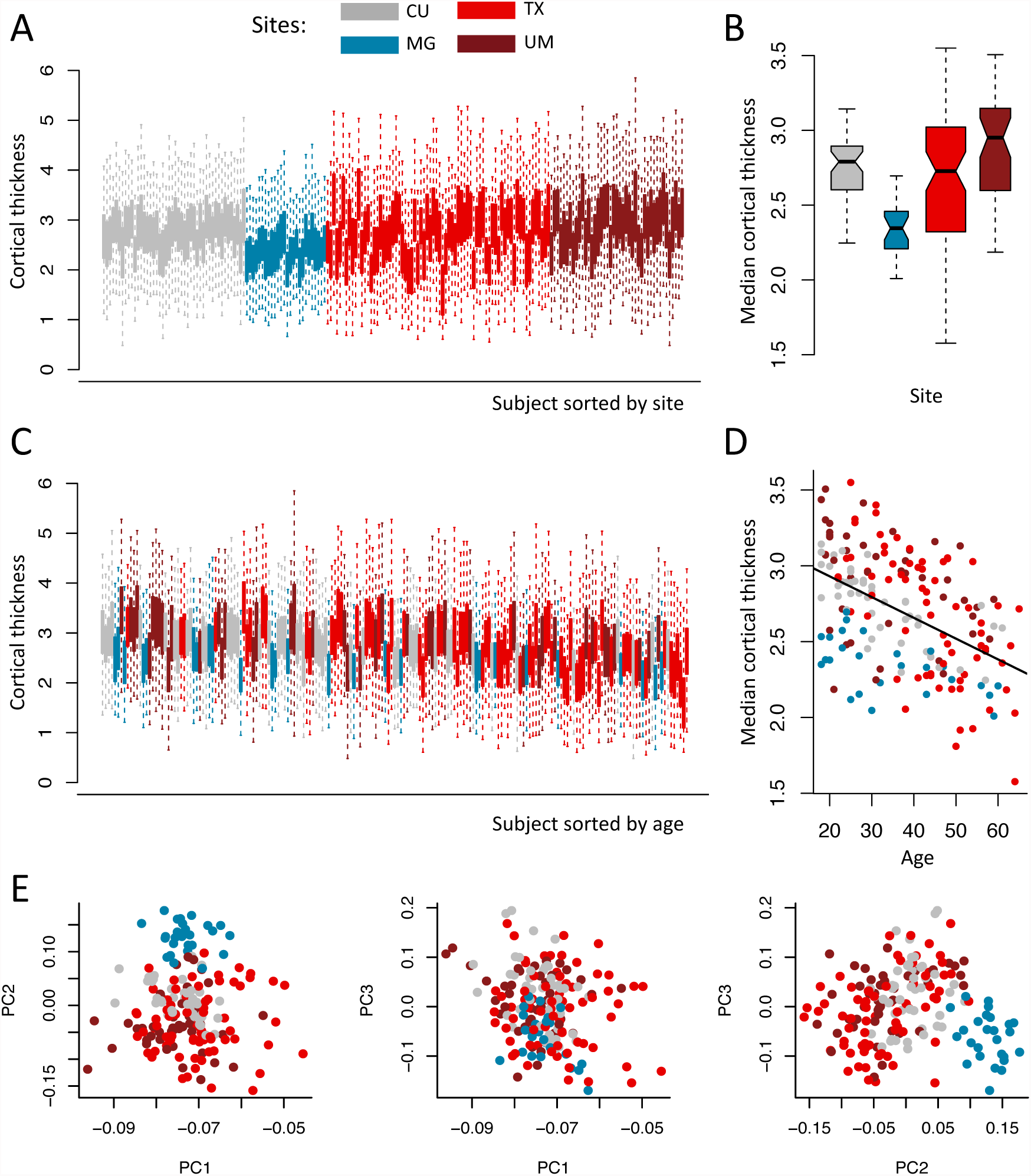
Visualization of sites effects in the EMBARC study. Plots are colored by imaging site: Columbia University (CU), University of Texas Southwestern (TX), Massachusetts General Hospital (MG) and University of Michigan (UM). (a) Boxplots of the cortical thickness sorted by site. Each boxplot represents the distribution of the 102 cortical regions for one subject. (b) Boxplots of the median cortical thickness, grouped by site. The MG site has lower median cortical thickness on average, while the TX and UM sites have higher variability. (c) Same as (a), but sorted by age. (d) Relationship between median cortical thickness and age, colored by site. (e) Plots of the first 3 principal components (PCs) from principal component analysis (PCA), colored by site. The second PC is highly associated with site.

In Figure 1c, the boxplots from Figure 1a were sorted by age. Consistent with the literature, we observe a global decrease of the cortical thickness measurements with age, and note that combining measurements from multiple sites adds variability to the trend (blue boxplots are shifted downwards). We also observe that the imaging sites are not distributed equally across the age span, with more younger subjects from the MG and CU sites (more blue and grey boxplots to the left) and older subjects coming from the TX site (more light red boxplots to the right). This indicates some counfouding level between imaging site and age. In Figure 1d, we present the median cortical thickness measurements as a function of age to visually inspect the global image-age relationship.

In Figure 1e, we present bivariate scatter plots of the first 3 principal components (PCs) from a principal component analysis (PCA) performed on the cortical thickness values. We note that the second PC is highly associated with site, confirming that a large proportion of the variation in the data is explained by site.

Finally, we present in Figure 2 the distribution of age, gender, HAMD score and QIDS score across imaging sites. This allows a visual inspection of potential confounding level between the different covariates and imaging site. The width of the boxplots represents the sample size at each site. We note that age is highly unbalanced across sites, with older subjects for the TX site, while the gender ratio seems to be equally distributed across sites. The QIDS score appears to be also unbalanced with respect to imaging site, and anti-correlated with age.

**Figure 2.**
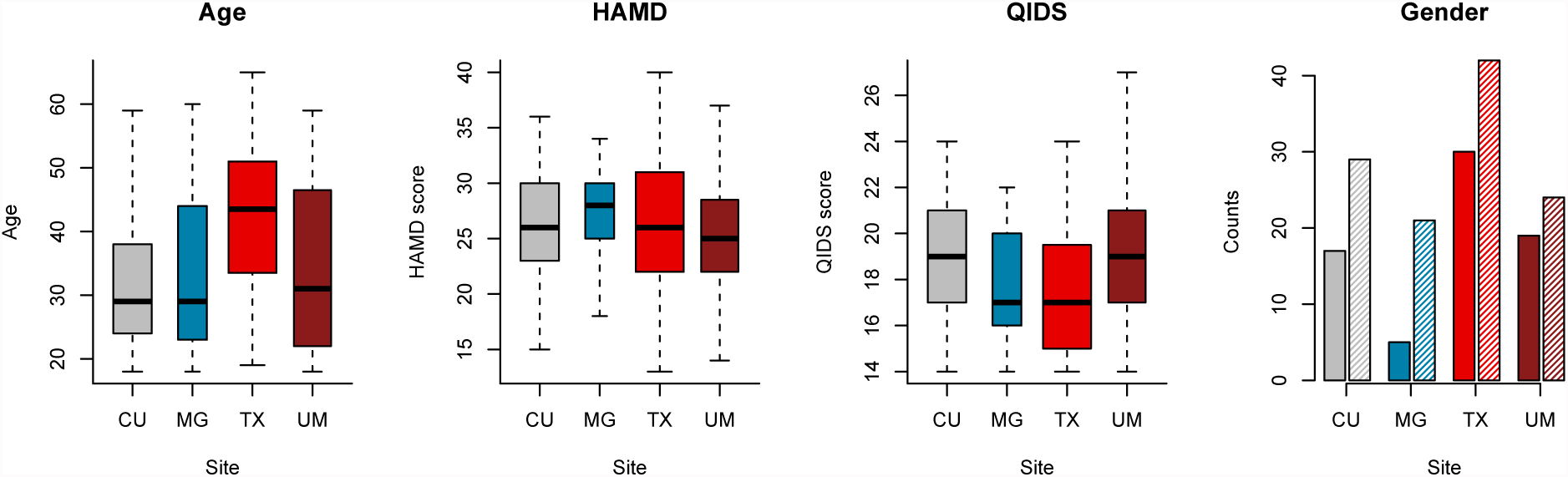
Covariates distribution in the EMBARC study. Distributions of age, gender, HAMD score and QIDS scores across sites for the EMBARC study. The width of the boxplots is proportional to the number of subjects scanned at each site. The full and shaded bars in the gender barplots represent males and females respectively. HAMD: Hamilton Depression Rating Scale; QIDS: Quick Inventory for Depression Symptomatology.

We also produced a similar visualization for the VDLC dataset in Figure A.1. We note that there is a clear positive shift in the cortical thickness measurements for images acquired on 3T scanners in comparison to images acquired on 1.5T scanners.

### 3.2 Quantification of site effects in the EMBARC study

We first evaluated the presence of global site effects in the cortical thickness measurements. For each subject, we summarized the measurements across regions by taking the median as a global measure. We present in Figure 1b the distribution of the median cortical thickness stratified by site. Using ANOVA, the median cortical thickness was significantly different across the four sites (p = 1.1 × 10^−10^). More specifically, the median cortical thickness for the MG site was significantly different from those of the three remaining sites, adjusting for multiple comparisons using Tukey’s method [Tukey, 1949].

Because the scale of the measurements can also be affected by scanner, we also compared the variances of the median cortical thickness measurement across sites. To do so, we performed the Bartlett’s sphericity test [Bartlett, 1937], which assesses whether or not the variances are homogeneous across sites. To avoid confounding of site with age and gender, we first regressed out the variation explained by age and gender; the test was significant (p = 1.8 x 10^−7^). We subsequently compared the pairwise site variances using the usual F-tests for variances ratio, and four of the pairs were significant after adjusting for multiple comparisons using Bonferroni correction: TX vs. CU, TX vs. MG, UM vs. CU, and UM vs. MG. The results are consistent with the spread of the boxplots presented in Figure 1b.

We then tested each ROI individually for site effects by calculating an ANOVA F-test. We obtained 53 ROIs significantly associated with site, using Bonferroni correction to adjust for multiple comparisons (adjusted *p* < 0.05). Because Bonferroni correction is a conservative approach to control for the family-wise error rate (FWER), we alternatively corrected for multiple comparisons using the permutation-based one-step maxT procedure [Westfall and Young, 1993, Dudoit et al., 2003], and obtained 60 ROIs significantly associated with site (adjusted *p* < 0.05, *B* = 10,000 permutations). We present in Figure B.1a the observed *R*
^2^ from ANOVA and the distribution of the maximum *R*
^2^ obtained from each permutation. To test for scanner-specific scaling effects, we also tested each feature individually for homogeneity of variances across sites using Bartlett’s test. We obtained 41 ROIs with variances significantly associated with site (adjusted *p* < 0.05, *B* = 10,000 permutations).

Quantification of the site effects for the VDLC study are provided in the Appendix A.1.

### 3.3 Removal of site effects with harmonization

To remove site effects in the EMBARC dataset, we applied three different harmonization techniques: (1) Residuals: removal of site effects estimated from linear regression; (2) Adjusted Residuals: removal of site effects estimated from linear regression, adjusting for biological covariates and (3) ComBat. In Figure 3, we show the empirical distributions of the site effects for both the location and scale parameters (dotted lines), together with the prior distributions estimated by ComBat (solid lines). We remind the reader that both the location and scale site effects are deviations from the grand mean. Consistent with the description of the site effects in the previous section, we note that the additive site effects (γ) are greater in magnitude for the MG site (Figure 3a, and the multiplicative site effects (δ) are greater than 1 on average for the TX and UM sites, and lower than 1 for the two remaining sites (Figure 3b). We note that the prior distributions fit the empirical distributions well for both the location and scale parameters; the ComBat procedure therefore appears appropriate for capturing these effects.

**Figure 3.**
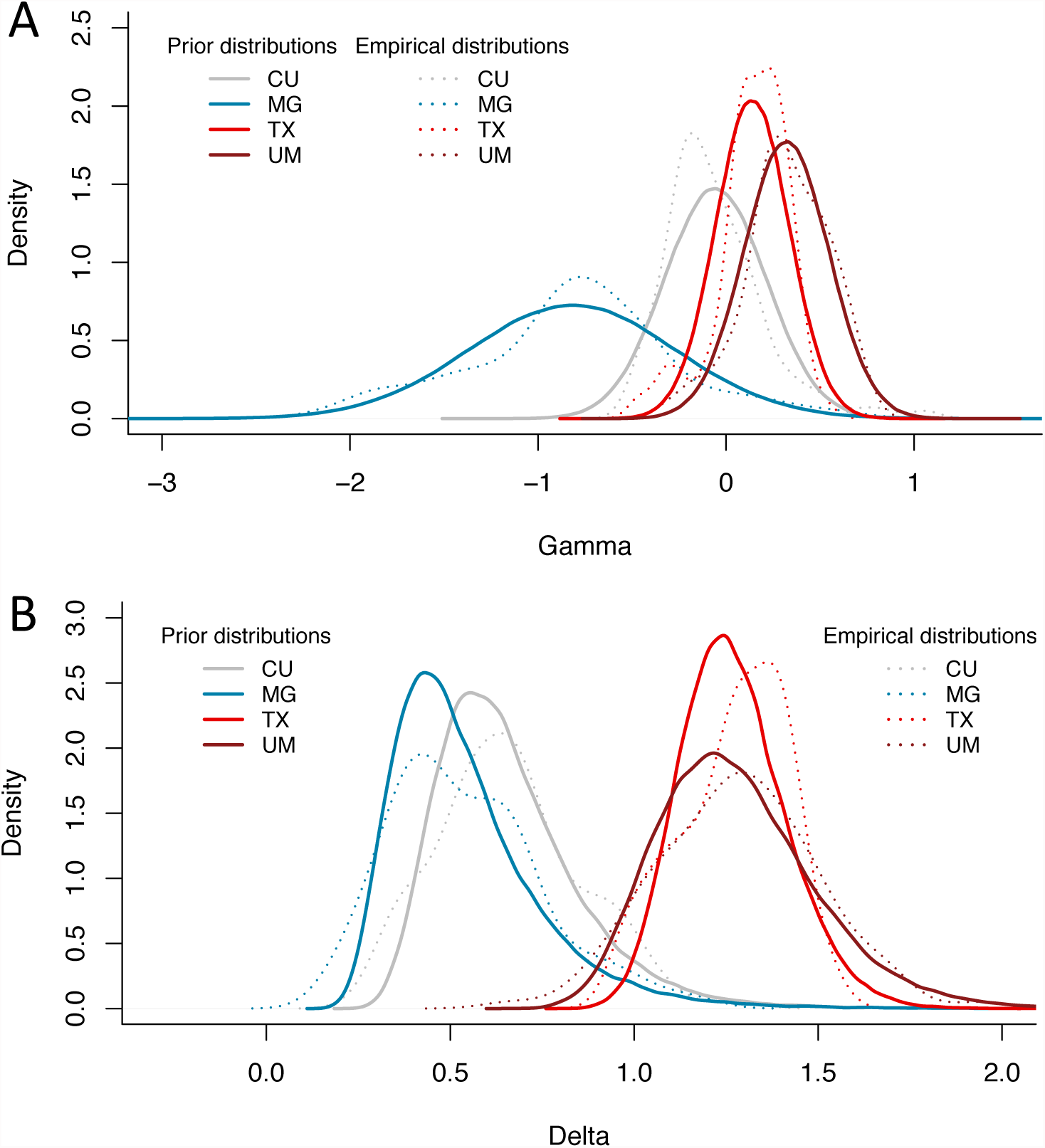
Prior distributions of the site effect parameters estimated by ComBat in the EMBARC study. Location and scale site-specific parameters estimated by ComBat, for the EMBARC study. (a) The ComBat-estimated prior distributions for the site-specific location parameters γ are shown in solid lines, and the empirical distributions of the site-specific location parameters are shown in dashed lines. (b) The ComBat-estimated prior distributions for the site-specific scale parameters δ are shown in solid lines, and the empirical distributions of the site-specific scale parameters are shown in dashed lines. The prior distributions fit well the empirical distributions for both the location and scale parameters.

To visualize whether or not most of the variation in the data was still associated with imaging site after harmonization, we first performed an unsupervised dimension reduction of the harmonized cortical thickness measurement using PCA. The data projected into the first two PCs are presented in the first column of Figure 4. We note that for all three harmonization methods, the data points appear to be distributed equally across sites. We also performed a linear discriminant analysis (LDA), a popular supervised dimension reduction that maximizes the projection coordinates to predict the data classes. Here, we use the imaging sites as the data classes to be predicted. We present the projected data in the second column of Figure 4. One can see that for the raw data, the data points cluster almost perfectly by imaging site. This is not surprising; all features are highly associated with site effects when not harmonized. After harmonization, site clusters are substantially attenuated.

**Figure 4.**
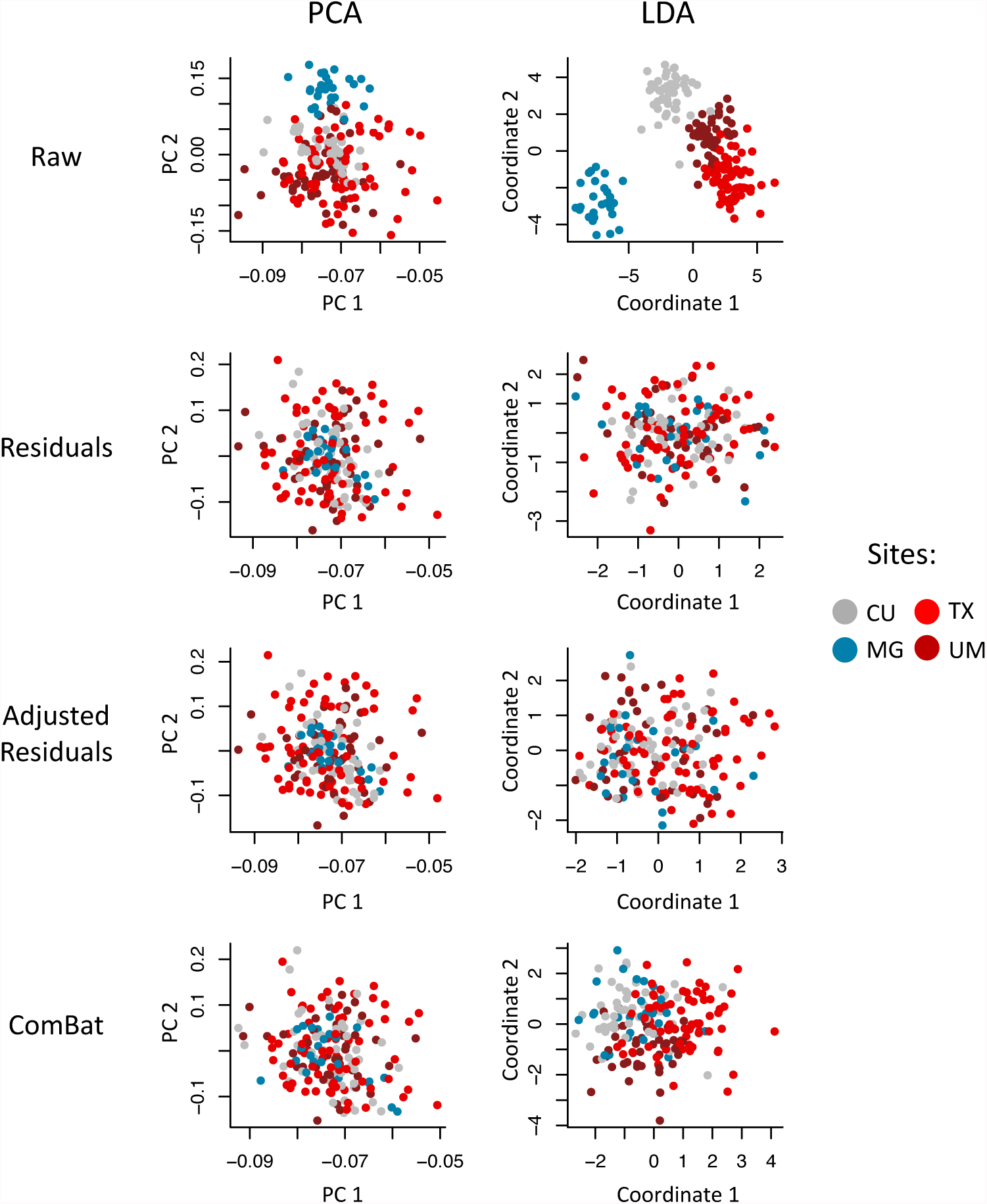
Supervised and unsupervised dimension reductions before and after harmonization for the EMBARC dataset. For each harmonization method, we first used principal component analysis (PCA) to reduce the dimension of the cortical thickness measurements in an unsupervised manner (agnostic of imaging sites). We present in the first column the projection of the data into the first two principal components (PCs) that explain most of the variation in the data. We also performed a supervised dimension reduction technique using linear discriminant analysis (LDA) using imaging site as a target variable. We present in the second column the projection of the data into the first two LDA coordinates. In both PCA and LCA, the first two coordinates are highly associated with site, while all harmonization methods removed most variation associated with site.

To formally test whether or not site effects remain after harmonization, we again used the different tests described in Section 3.2. Using ANOVA F-tests, all methods corrected for mean site differences in the median cortical thickness: *p* = 0.997 for Residuals, *p* = 0.0498 for Adjusted Residuals and *p* = 0.0473 for ComBat. We also tested for site-specific scaling effect in the measurements using Bartlett’s sphericity test. We found that only ComBat was able to remove the scaling effects associated with site (*p* = 0.42). The site-specific variances remained largely uncorrected for both the Residuals (*p* = 2.53 x 10^−8^) and Adjusted Residuals (*p* = 3.08 x 10^−8^) methods. This is not surprising; only the ComBat harmonization method is able to model scaling factors associated with site. We also tested each ROI individually for remaining site effects. For all harmonization methods, none of the ROIs was significantly associated with site, using either the Bonferroni or the maxT adjustment.

Finally, to further investigate if site effects were entirely removed for each of the harmonization method, we attempted to predict imaging site from the harmonized cortical thickness features. More specifically, we used the support vector machine (SVM) [Cortes and Vapnik, 1995] classification algorithm, with radial basis kernel, to predict site from the imaging features. The SVM is largely used in the imaging community in the context of multivariate pattern analysis (MPVA) for understanding and discovering patterns associated with a disease outcome, for instance. A harmonization method that is successful in removing site effects will result in a lower SVM accuracy when attempting to predict site. Using *B* = 10,000 repetitions of a 10-fold cross-validation, we estimated an average accuracy for each method. For the raw values, the SVM prediction achieved an average of 76.6% classification accuracy. For the residuals and adjusted residuals methods, the average accuracies were 40.5% and 38.7% respectively. The ComBat method resulted in the lower average accuracy (36.3%). Using a permutation-based approach to generate a null distribution (B = 10,000), a SVM classification by chance attained on average 36.9% accuracy. This indicates the Adjusted Residuals and ComBat were best for the removal of site effects in the cortical thickness measurements. In comparison to the adjusted residuals, we note that the ComBat method additionally removes site-specific scaling effects. This could explain the better performance in the SVM, in which the covariance structure is implicitly used for predicting the class labels.

### 3.4 Associations with age

While it is important to show that a harmonization method successfully removes site effects, it is equally important to show that the method preserves the biological variability in the data; a method that removes both site effects and biological effects has no scientific use. To investigate whether or not the different harmonizations presented in this paper perform well at preserving biological variability, we use age as a variable of interest. Again, we use the EMBARC dataset to demonstrate the main results.

We first assessed the proportion of variation explained by age before and after harmonization. Without harmonization, the percentage of variation in the average cortical thickness explained by age was 23%. This was calculated using the usual coefficient of variation *R*
^2^ from linear regression with median cortical thickness as the outcome. For the unadjusted Residuals method, this percentage was increased to 26%, and for both the Adjusted Residuals and ComBat, the percentage was increased to 33%. The fact that the Unadjusted Residuals did not substantially increase the association with age is not surprising; we observed that age was confounded with imaging site, and therefore removing site effects without adjusting for age will also remove variation in the imaging features associated with age. On the other hand, both the Adjusted Residuals and ComBat strengthened the expected inverse relationship between age and cortical thickness by removing site effects, but also by preserving biological variability in the data.

We also evaluated the effects of harmonization on the prediction of age using the harmonized cortical thickness measurements. For prediction, we used two different algorithms: linear regression, and the popular support vector regression (SVR) algorithm, also commonly called *ϵ*-SVM regression, using a radial kernel. The *ϵ-*SVM regression paradigm is similar to the regular SVM, but for a continuous outcome. For each algorithm, we used the harmonized cortical thickness measurements as imaging features inputs X to predict age. For each harmonization method, we randomly partitioned the subjects into k folds, and trained the prediction algorithm on *k –* 1 folds. We then predicted the age of the remaining subjects (testing dataset) and calculated the root-mean-square error (RMSE). We repeated the random sampling *B* = 1000 times, for *k* ϵ {3,5,10}, to obtain a distribution of the RMSE for each method at each *k*.

In Figure 5, we present the results from linear regression. For the three values of *k*, we observe that the data harmonized with the unadjusted Residuals do not perform well, while both the Adjusted Residuals and ComBat improve the prediction of age upon the raw data. In Figure 6, we present the results from *ϵ*-SVM regression. For the three values of *k*, we observe that the data harmonized with the unadjusted Residuals do not perform well, while the Adjusted Residuals and ComBat perform very similarly to the raw data. Interestingly, we note that while the Adjusted Residuals and ComBat methods improve the prediction in the case of linear regression, very little improvement was obtained in *ϵ*-SVM regression. One explanation could be that the different harmonization methods presented in this paper are linear models, and therefore are optimal for prediction using linear prediction, but not necessarily for the *ϵ*-SVM regression.

**Figure 5.**
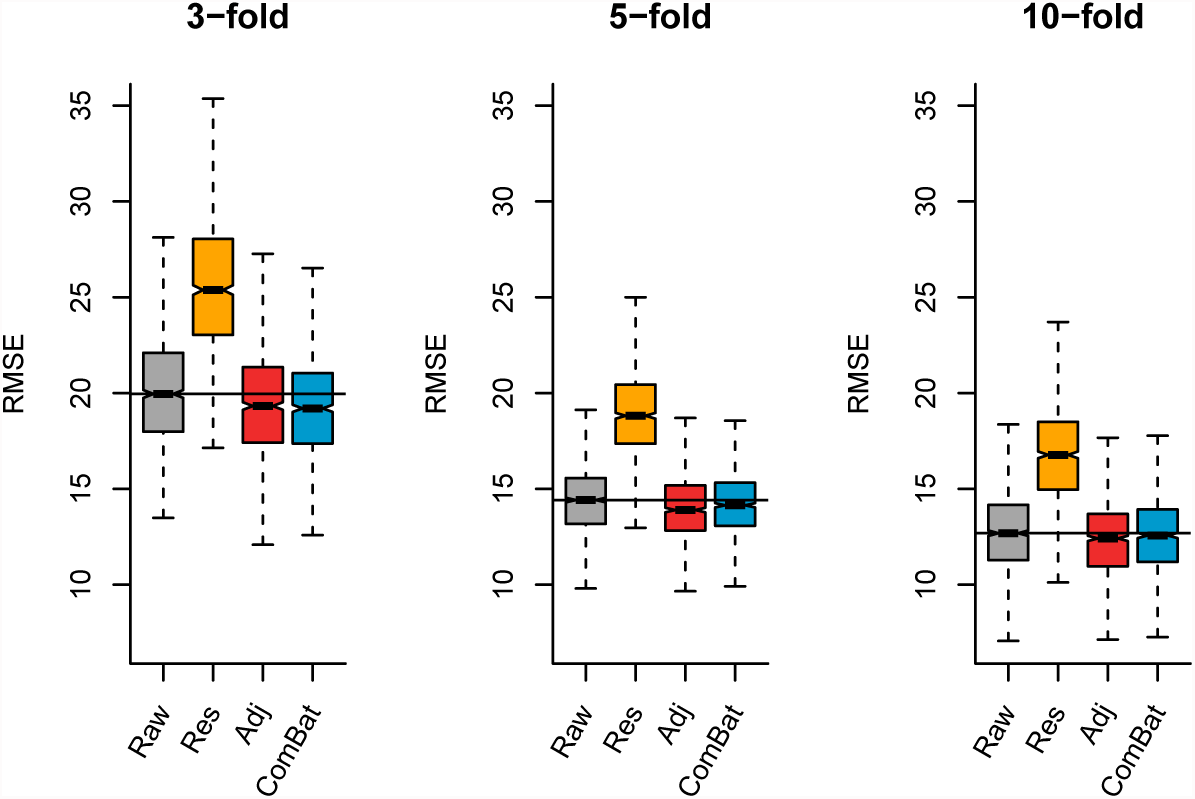
Root-mean-square error (RMSE) for age prediction using linear regression. Using *k*-fold validation for *k* ∈ {3,5,10} for *B* = 1000 random samplings, we calculated the RMSE on a testing dataset for the predicted age using linear regression. For the different harmonization methods, we used the harmonized cortical thickness measurements as input image features to train the algorithm.

**Figure 6.**
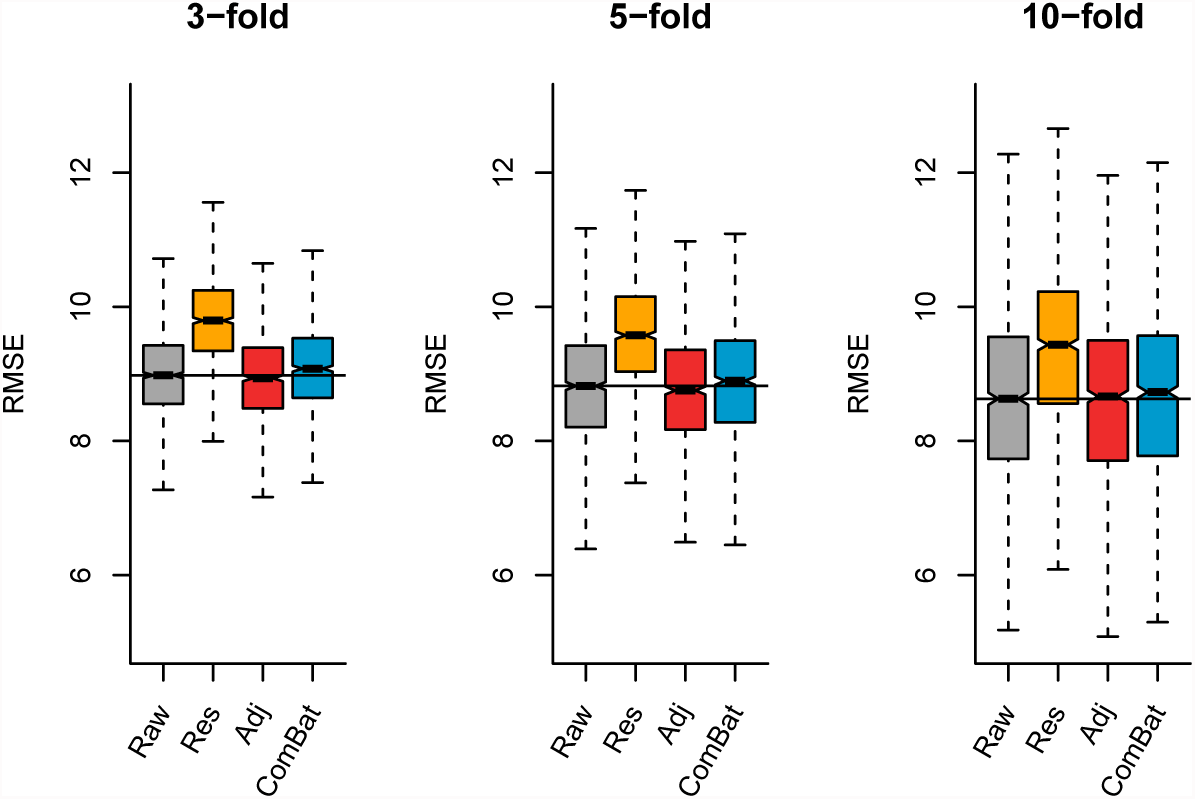
Root-mean-square error (RMSE) for age prediction using *ϵ*-SVM. Using *k*-fold validation for *k* ∈ {3,5,10} for *B* = 1000 random samplings, we calculated the RMSE on a testing dataset for the predicted age using *ϵ*-SVM. For the different harmonization methods, we used the harmonized cortical thickness measurements as input image features to train the algorithm.

### 3.5 Life-span study by harmonizing the EMBARC and VDLC datasets

While the two studies present in this paper have a different age range ([18,65] y.o. for the EMBARC study; [58,95] y.o. for the Vascular study), there is some overlap between the two age ranges (see Figure 7, first panel). For the study of life-span trajectories, it is sometimes necessary to combine data from multiple studies, with each individual study often targeting participants from a specific age range. We show here that even though different scanners were used across the studies, it is possible to combine and harmonize the data, to remove the scanner effects, and thereby improve the correlation between the imaging outcome and biological factors of interest, namely age.

**Figure 7.**
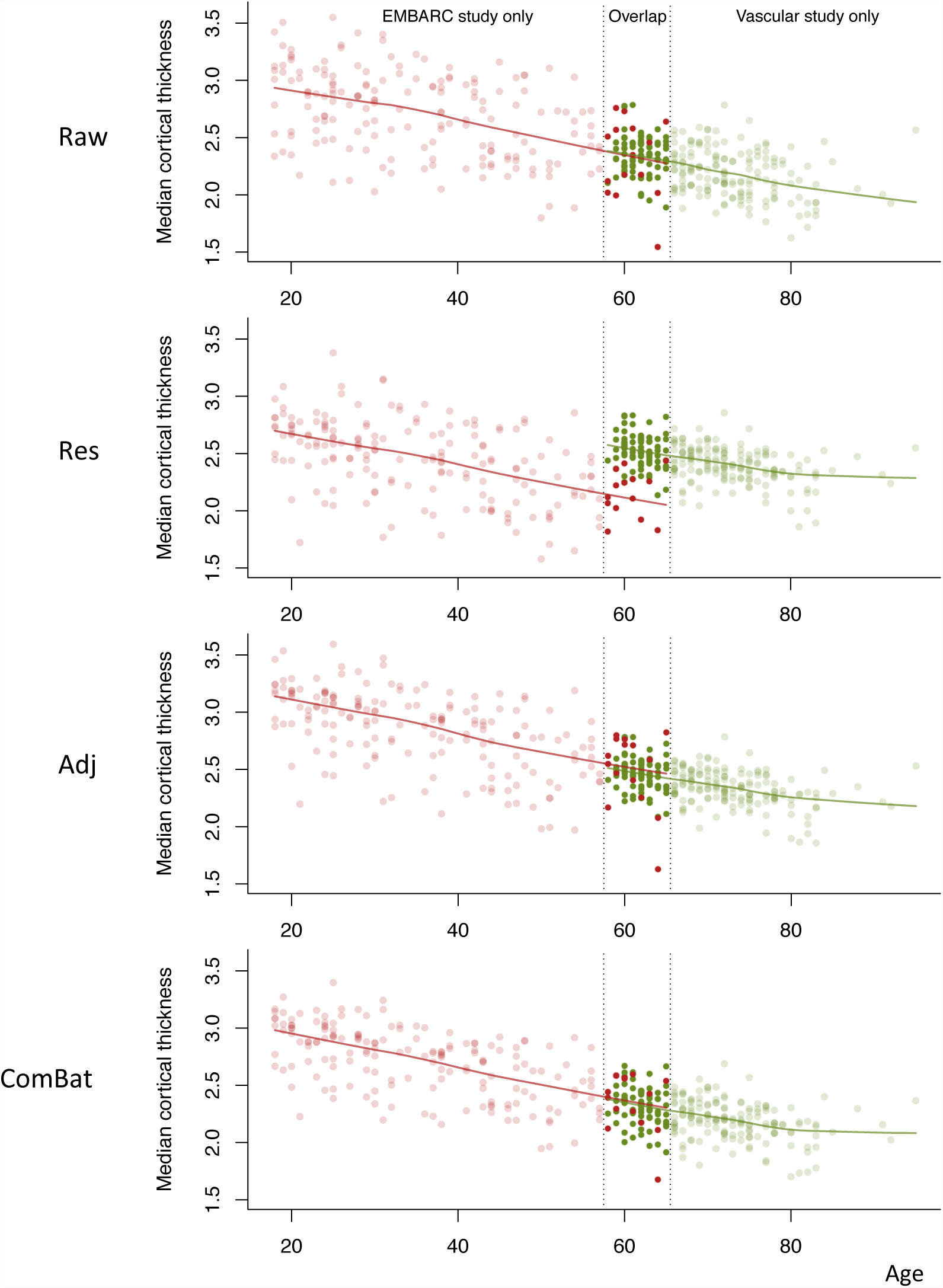
Median age trajectory before and after harmonization. The EMBARC and Vascular studies were combined using different harmonizations. The red dots represent the median cortical thickness for the EMBARC study participants, and the greens dots represent the median cortical thickness for the Vascular study participants. The curves represent the lowess fitted values for each study separately.

We present the relationship between median cortical thickness and age, before and after harmonization in Figure 7 with data points colored by study (red for EMBARC and green for Vascular). One can observe an overlap in the age span between the two studies, and that inter-subject variation seems to be higher in the EMBARC subjects in the raw data. This can be explained by the large variation between the four scanners in the EMBARC, as discussed previously in the Results section. For each method, we calculated the correlation between the median cortical thickness and age. For the unharmonized data, we obtained a correlation of −0.70. For the unadjusted Residuals, we obtained a correlation of −0.26. Such a weaker correlation is not surprising; both studies have a vastly different age range, and therefore blindly harmonizing the data for site without adjusting by age will diminish the age effect across the life span. For the Adjusted Residuals, we obtained a correlation of −0.77, and we obtained a correlation of −0.79 for ComBat. Both adjusted residualization and ComBat were effective at decreasing the inter-subject variability by removing scanner effects, while preserving the trend associated with age across the life-span.

## 4 Discussion

With the increasing complexity of study design in multi-site neuroimaging studies, the neuroscience community needs robust, validated, and computationally feasible methods for addressing the critical impact of non-biological sources of data variation. We use the term “harmonization” to refer to the process of combining data from multiple sites and removing the unwanted variability associated with scanner.

In this paper, we proposed to use the ComBat algorithm, previously developed to deal with batch effects in the study of gene expression data, as a reliable harmonization method for combining cortical thickness measurements across sites. This was motivated by its previously documented excellent performance for harmonizing voxel-wise fractional anisotropy (FA) and mean diffusivity (MD) measurements [Fortin et al., 2017], two common DTI scalar maps. Using two large multi-site studies, EMBARC and VDLC, we presented a general approach for identifying unwanted sources of variance in neuroimaging data using exploratory, univariate, and multivariate tests for scanner effects. We then showed that ComBat is effective at removing nuisance variability associated with scanners, while preserving the age effects in the cortical thickness across participants. We also showed that ComBat can be used to combine those two large studies, with a vastly different age range, to study cortical thickness across the life span.

We compared the ComBat harmonization algorithm to two commonly used scanner effect correction methods: residualization and adjusted residualization. The latter method adjusts for covariates of interest (for instance age) in the removal of site effects. ComBat is similar to the adjusted method, except that it additionally models scanner-specific scaling effects. ComBat also uses a Bayesian framework to improve the stability of the estimated parameters in small sample sizes. ComBat is easy to apply and has minimal computational overhead. Equally importantly, we have developed open-source, easy-to-use code for applying this algorithm in R, Matlab, and Python. This ensures that the ComBat algorithm can be seamlessly integrated into any existing processing pipelines.

We note that several other harmonization techniques have been previously proposed in the context of other imaging modalities. For conventional MRI studies, intensity normalization techniques have been developed to make the image intensities comparable across studies, including histogram matching [Nyúl et al., 2000], WhiteStripe [Shinohara et al., 2014] and RAVEL [Fortin et al., 2016]. Another method, called source-based morphometry, uses independent component analysis (ICA) to remove variability associated with certain scanner parameters in structural MRI [Chen et al., 2014]. For diffusion tensor imaging (DTI) studies, it has been proposed to use spherical harmonics to harmonize data across studies, using a reference site to create pairwise site transformations [Mirzaalian et al., 2016]. It has also been proposed to use functional normalization, originally developed in [Fortin et al., 2014], for harmonizing DTI scalar maps. We note that the aforementioned harmonization techniques could not be readily applied to cortical thickness.

In the future, we plan to develop a time-dependent ComBat algorithm for understanding scenarios where subjects were scanned over multiple time points, and for which scans were acquired on different scanners, or on the same scanners but with different scanning parameters. We are also planning on improving the performance of ComBat in the presence of confounding by implementing an inverse probability weighting (IPW) scheme into the algorithm. IPW has been shown to improve prediction when the outcome of interest is confounded with another covariate [Linn et al., 2016]. This has the potential to improve the performance of ComBat for age prediction using the SVM regression framework, as well as for other prediction methods.

## 5 Software

All postprocessing analysis was performed in the R statistical software (version 3.2.0). For ComBat, the reference implementations from the sva package was used. All figures were generated in R with customized and reproducible scripts. We have adapted and implemented the ComBat methodology to imaging data, and the software is available in R and Matlab (https://github.com/Jfortin1/ComBatHarmonization) and in Python (https://github.com/ncullen93/neuroCombat).

## Abbreviations

ANTs: Advanced normalization tools
AD: Alzheimer’s disease
ADNI: Alzheimer’s Disease Neuroimaging Initiative
ANOVA: Analysis of variance
DTI: Diffusion tensor imaging
EB: Empirical Bayes
EMBARC: Establishing Moderators and Biosignatures of Antidepressant Response in Clinical care
FA: Fractional anisotropy
FWER: Family-wise error rate
GPR: Gaussian process regression
HAMD: Hamilton Depression Rating Scale
IPW: Inverse probability weighting
LDA: Linear discriminant analysis
LLD: Late-life depression
MASQ: Mood and Anxiety Symptom Questionnaire
MCI: Mild cognitive impairment
MD: Mean diffusivity
MVPA: Multivariate pattern analysis
OASIS: Open Access Series of Imaging Studies
PC: Principal component
PCA: Principal component analysis
QIDS: Quick Inventory for Depression Symptomatology
RMSE: Root-mean-square error
ROI: Region of interest
STAI: Spielberger State-Trait Anxiety Inventory
SVM: Support vector machine
SVR: Support vector regression
VDLC: Vascular disease: Longitudinal changes

## Authors contributions

JPF developed the methodology. WDT, PA, CC, MF, PJM, MM, RVP, MLP, MHT and MMW recruited the participants and acquired the data. IA processed the data. JPF and NC developed software and analyzed the data. JPF, NC, RTS and YIS wrote the manuscript. RTS and YIS supervised the work. All authors read and approved the final manuscript.

## Acknowledgements

The research was supported in part by R01NS085211 and R21NS093349 from the National Institute of Neurological Disorders and Stroke, R01MH112847 from the National Institute of Mental Health. The EMBARC study was supported by U01MH092221 and U01MH092250. The VDLC study was supported by R01MH060697, R01MH074916, R01MH078216 and NCT00045773. The content is solely the responsibility of the authors and does not necessarily represent the official views of the funding agencies.

## Appendix A

### A.1 Quantification of site effects in the VDLC study

We present in Figure A.1b the distribution of the median cortical thickness stratified by scanner. Using ANOVA, the median cortical thickness was significantly different across the seven scanners (p = 2.2 × 10^−16^). Not surprisingly, the median cortical thicknesses from each of the 3T scanners were significantly different from those of each of the 1.5T scanner, adjusting for multiple comparisons using Tukey’s method.

We also compared the variances of the median cortical thickness measurement across scanners. To do so, we performed the Bartlett’s sphericity test, which estimates whether or not the variances are homogeneous across scanners. To avoid confounding of scanner with age and gender, we first regressed out the variation explained by age and gender; the test was significant (p = 0.0013).

We then tested each ROI individually for site effects by calculating an ANOVA F-test. We obtained 86 ROIs significantly associated with site, using Bonferroni correction to adjust for multiple comparisons (adjusted *p* < 0.05), and 87 ROIs using the permutation-based one step maxT procedure (adjusted *p* < 0.05, *B* = 10,000 permutations). We present in Figure B.1b the observed *R*
^2^ from ANOVA and the distribution of the maximum *R*
^2^ obtained from each permutation. To test for scanner-specific scaling effects, we also tested each feature individually for homogeneity of variances across sites using Bartlett’s test. We obtained 4 ROIs with variances significantly associated with site (adjusted *p* < 0.05, *B* = 10,000 permutations).

### A.2 Removal of scanner effects in the VDLC study with harmonization

In Figure A.2, we show the empirical distributions of the site effects for both the location and scale parameters (dotted lines), together with the prior distributions estimated by ComBat (solid lines). We remind the reader that both the location and scale scanner effects are deviations from the grand mean. Consistent with the description of the site effects in the previous section, we note that the additive scanner effects (*γ*) are greater in magnitude for the 3T scanners. The multiplicative scanner effects (*δ*) are shown in Figure A.2b. We note that the prior distributions fit the empirical distributions well for both the location and scale parameters; the ComBat procedure therefore appears appropriate for capturing these effects.

To visualize whether or not most of the variation in the data was still associated with scanner after harmonization, we first performed an unsupervised dimension reduction of the harmonized cortical thickness measurement using PCA. The data projected into the first two PCs are presented in the first column of Figure A.3. We note that for all three harmonization methods, the data points appear to be distributed equally across scanners. We also performed LDA using scanners as the data classes. We present the projected data in the second column of Figure A.3. One can see that for the raw data, there is a clear separation between the different types of scanners. Interestingly, the data from the D SIGNA scanner appear to cluster separately; We note that this is the only GE scanner in the VDLC study. After harmonization, clusters associated with scanner are substantially attenuated.

**Figure A.1.**
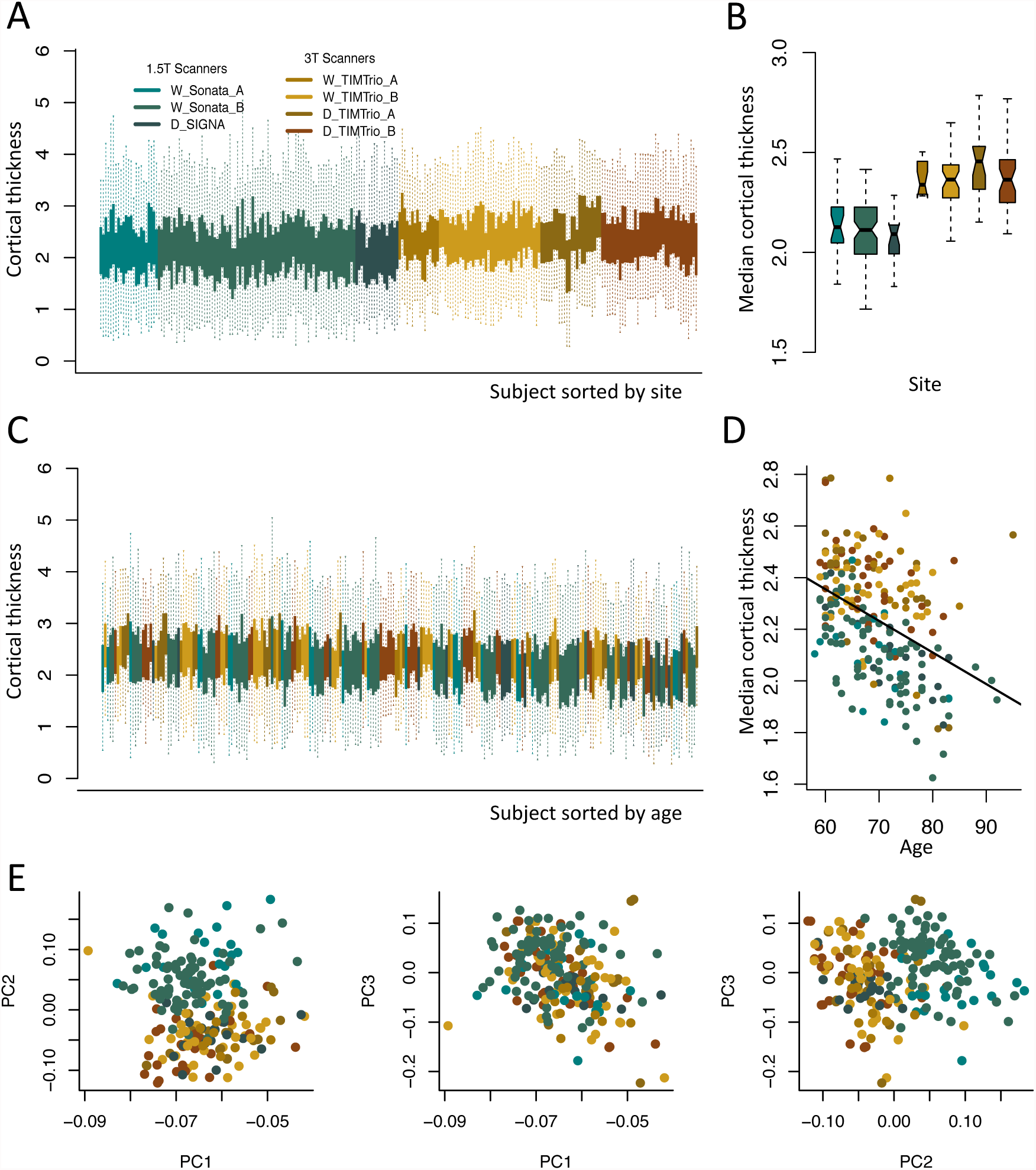
Visualization of sites effects in the Vascular study. Plots are colored by scanner. The green shades represent the 1.5T scanners, while the brown shades represent the 3T scanners. (a) Boxplots of the cortical thickness sorted by site. Each boxplot represents the distribution of the 98 cortical regions for one subject. (b) Boxplots of the median cortical thickness, grouped by scanner. The measurements derived from 1.5T scanners are substantially lower than measurements from 3T scanners. (c) Same as (a), but sorted by age. (d) Relationship between median cortical thickness and age, colored by scanner. (e) Plots of the first 3 principal components (PCs) from principal component analysis (PCA), colored by scanner. The second PC is highly associated with scanner.

**Figure A.2.**
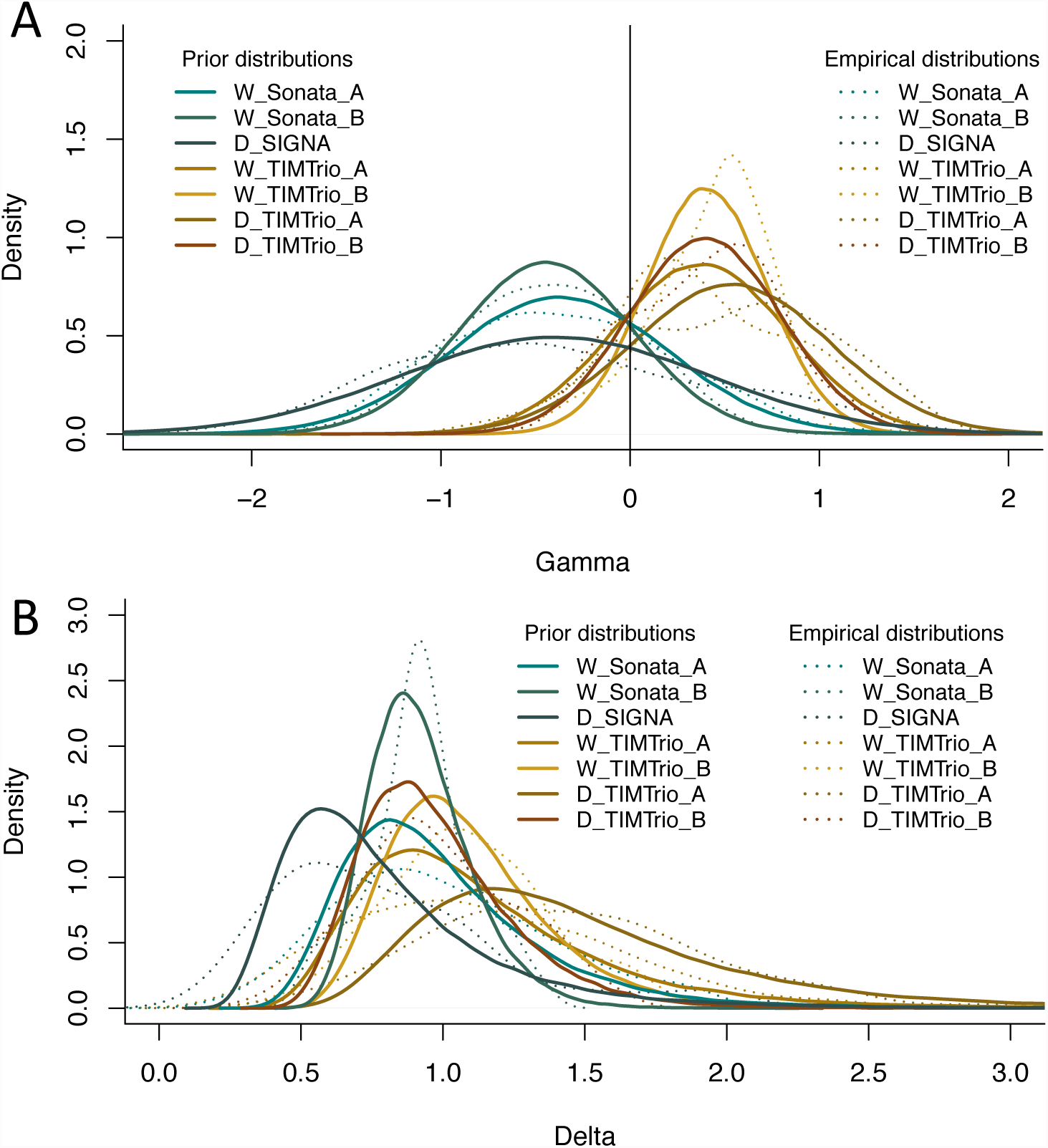
Prior distributions of the site effect parameters estimated by ComBat in the Vascular study. Location and scale site-specific parameters estimated by ComBat, for the EMBARC study. (a) The ComBat-estimated prior distributions for the site-specific location parameters γ are shown in solid lines, and the empirical distributions of the site-specific location parameters are shown in dashed lines. (b) The ComBat-estimated prior distributions for the site-specific scale parameters δ are shown in solid lines, and the empirical distributions of the site-specific scale parameters are shown in dashed lines. The prior distributions fit well the empirical distributions for both the location and scale parameters.

**Figure A.3.**
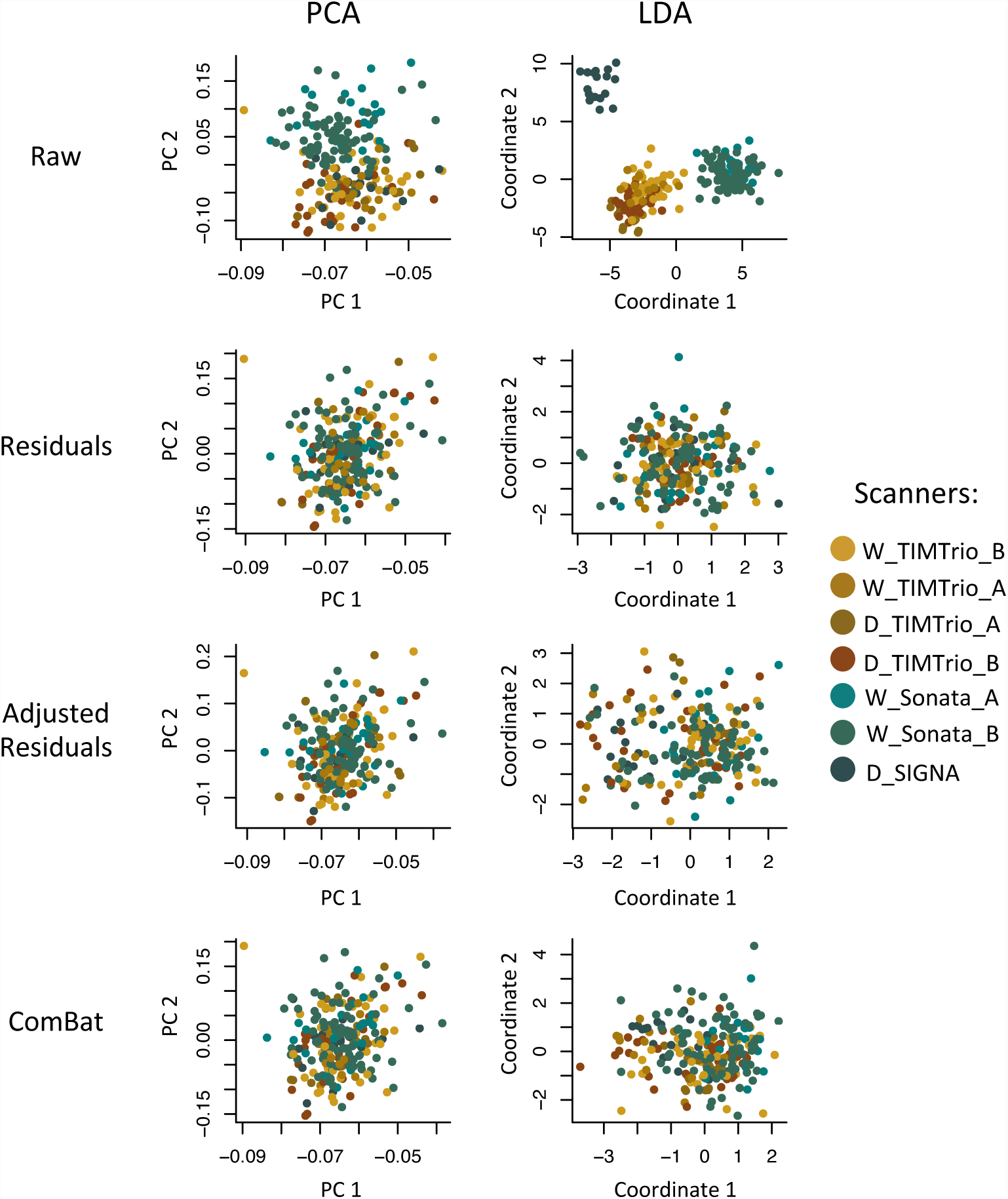
Supervised and unsupervised dimension reductions before and after harmonization for the Vascular dataset. For each harmonization method, we first used principal component analysis (PCA) to reduce the dimension of the cortical thickness measurements in an unsupervised manner (agnostic of imaging sites). We present in the first column the projection of the data into the first two principal components (PCs) that explain most of the variation in the data. We also performed a supervised dimension reduction technique using linear discriminant analysis (LDA) using imaging site as a target variable. We present in the second column the projection of the data into the first two LDA coordinates. In both PCA and LCA, the first two coordinates are highly associated with site, while all harmonization methods removed most variation associated with site.

Using ANOVA F-tests, all methods corrected for mean scanner differences in the median cortical thickness: *p* = 0.99 for Residuals, *p* = 0.94 for Adjusted Residuals and *p* = 0.94 for ComBat. We also tested for scanner-specific scaling effects in the measurements using Bartlett’s sphericity test. We found that only ComBat was able to remove the scaling effects associated with scanner (*p* = 0.46). Scanner-specific variances remained present in both the Residuals (*p* = 0.03) and Adjusted Residuals (*p* = 0.01) methods. Finally, we tested each ROI individually for remaining scanner effects. For all harmonization methods, none of the ROIs was significantly associated with scanner, using either the Bonferroni or the *maxT* adjustment.

As for the EMBARC study, we used the SVM with radial basis kernel, to assess prediction of scanner from the imaging features. Again, a harmonization method that is successful in removing scanner effects will result in a lower SVM accuracy when attempting to predict scanner. Using *B* = 10,000 repetitions of a 10-fold cross-validation, we estimated an average accuracy for each method. For the raw values, the SVM prediction achieved an average of 67.7% classification accuracy. For the residuals and adjusted residuals methods, the average accuracies were 43.4% and 44.4% respectively. The ComBat method resulted in the lower average accuracy (41.0%).

## Appendix B

**Figure B.1.**
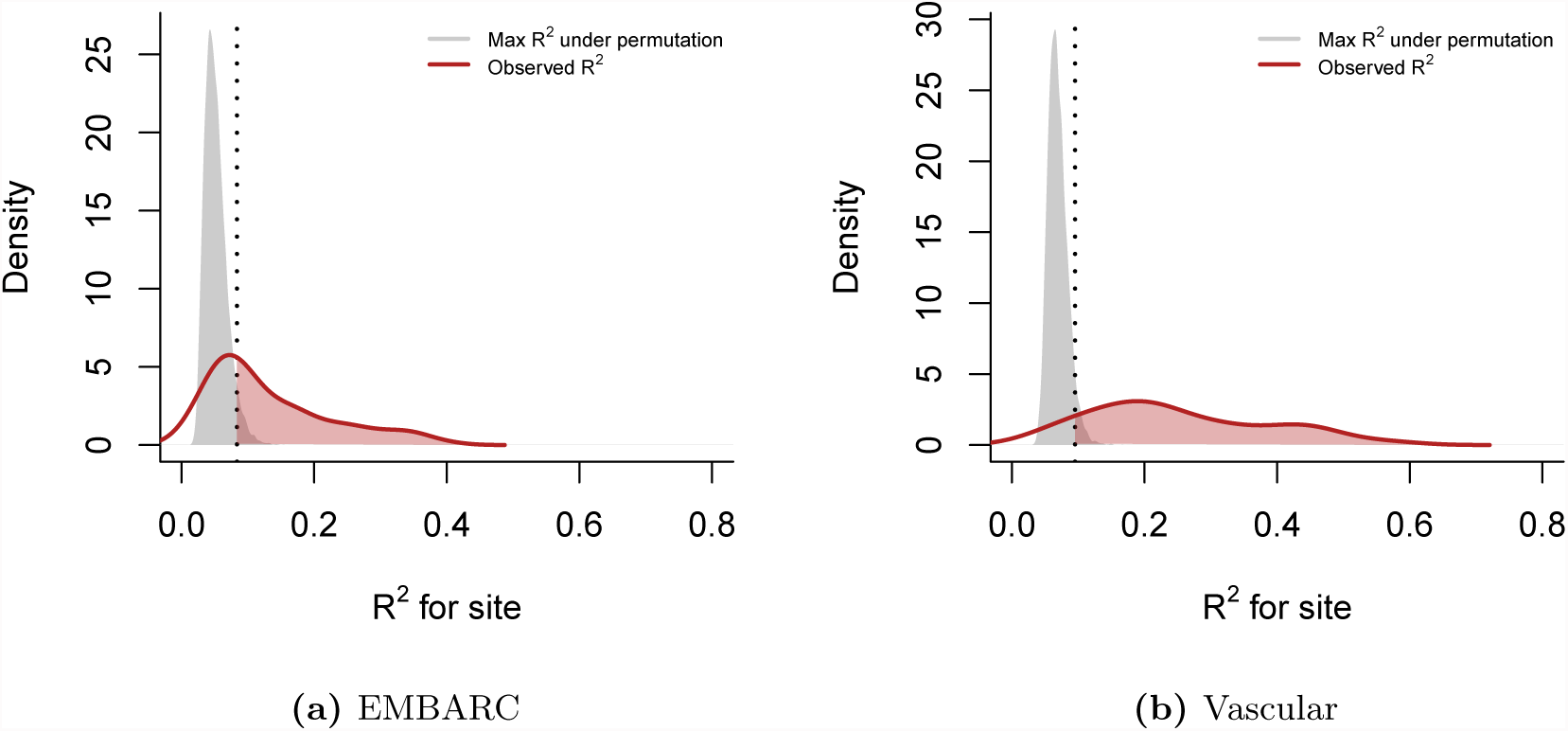
Variance explained by imaging site (*R*
^2^). For each feature, we calculated the coefficient of determination *R*
^2^ between cortical thickness and imaging site. We present the densities of *R*
^2^ (red lines) for the (a) EMBARC study and the (b) Vascular study. To obtain a measure of significance and to correct for multiple comparisons, we performed a one-step max *R*
^2^ procedure. Briefly, we permuted the site labels *B* = 10,000 times, recalculated the *R*
^2^ values and retained the maximum *R*
^2^ value at each permeation. The grey densities represent the distribution of the maximum *R*
^2^’s. The vertical dashed line indicates the 95% quantile of the maximum *R*
^2^ distribution. The features above that threshold are significant at the *α* = 0.05 level (shaded red area). Most features remained significant after adjustment.

